# Task-Dependent Effects of Channel Density on Multivariate Pattern Analysis and Spatial Representational Similarity Analysis: Evidence from Three EEG Datasets

**DOI:** 10.64898/2026.01.26.701731

**Authors:** Limin Hou, Guanghui Zhang, Yuxing Hao, Tommi Kärkkäinen, Fengyu Cong

## Abstract

Despite the widespread use of EEG decoding and representational similarity analysis (RSA), the extent of the impact of electrode density on these multivariate approaches has not been systematically evaluated, particularly across different cognitive domains. Here, we systematically compared four electrode configurations (19, 32, 64, and 128 channels) across three EEG datasets encompassing visual perception, visual search, and emotional processing. All datasets were recorded at 128 channels and subsequently reduced to 64, 32, and 19 channels via electrode subset selection based on the international 10-20 system. Time-resolved decoding and spatial RSA were computed for each configuration with paired statistical evaluation. Decoding sensitivity demonstrated clear task dependence: in the color-harmony paradigm, performance improved markedly with higher densities, consistent with spatially distributed perceptual representations, whereas decoding in the icon-search and dot-probe tasks remained stable across densities, indicating diminishing returns of additional electrodes although all configuration yielded robust decoding accuracies that above chance level. RSA results were comparatively robust, with relative representational similarity structures largely preserved even for 16 channel counts. These findings demonstrate that low-density setups can be sufficient, offering practical guidance for EEG channel selection and supporting the use of low-density EEG in real-world applications, including wearable systems, mobile EEG recordings, and clinical or rehabilitation settings.

## 1 Introduction

Electroencephalography (EEG) is a non-invasive technique for recording electrical activity from the brain. It is essential to cognitive neuroscience because of its high temporal resolution, ease of use, and scalability (Cohen, 2014). In recent years, the growing adoption of high-density EEG systems has enabled researchers to analyze the spatiotemporal dynamics of brain activity with increasing precision (Cutini et al., 2021; Song et al., 2015). However, in practical applications, high-density EEG equipment remains costly and time-consuming to set up (Di Flumeri et al., 2019; Kleffner-Canucci et al., 2012). Furthermore, in specific scenarios such as long-term monitoring, portable EEG use, or resource-constrained research environments, researchers often need to work with few channel configurations, for example using 32, 19, or even fewer electrodes (Johnson et al., 2011; M. Wang et al., 2021).

In recent years, multivariate pattern analysis (MVPA) has become a widely used approach in cognitive neuroscience for decoding time-resolved neural representations from EEG, capturing subtle and more discriminative patterns of neural activity than univariate approaches (Grootswagers et al., 2017; Zhang, Wang, et al., 2025; Zhang & Luck, 2025). In EEG MVPA analyses that are based on sensor-level spatial patterns across electrodes at each time point, decoding performance can be affected by the spatial sampling density of the electrode array (Gurariy et al., 2022; Robinson et al., 2017). Previous studies have extensively examined the influence of EEG channel density on signal analysis outcomes, including its effects on spatial resolution (Corley & Huang, 2018; Srinivasan, 1999; Vaisanen, 2008), feature extraction (Asayesh et al., 2024), and decoding performance (Huang et al., 2026; Nogueira, Cosatti, et al., 2019). Most research has found that classification accuracy decreases with the number of channels decreases (Lee et al., 2022; Nogueira, Dolhopiatenko, et al., 2019) and few channel numbers can impair the spatial resolution of EEG signals (Allouch et al., 2023; Ryynanen et al., 2004). In addition, studies using MVPA or neural decoding approaches have consistently shown that high-density EEG systems tend to yield better classification or decoding performance (M. Wang et al., 2021). However, recent studies have demonstrated that even with only 16 electrodes, it is possible to successfully decode visual stripe features and category information of complex natural images, indicating that low-density EEG can also provide robust neural representations under specific conditions (Huang et al., 2026). These findings challenge the common assumption that “more channels are always better,” yet it remains unclear whether such robustness consistently across different cognitive tasks, such as perception, memory, or decision-making.

Furthermore, research on the impact of channel count on representational similarity analysis (RSA) remains relatively limited. RSA is a multivariate method that quantifies the similarity of neural activity patterns to reveal the representational structure of information in the brain (Kriegeskorte et al., 2008). In time-resolved EEG studies, RSA is typically implemented by extracting, at each time point, a spatial activity vector across all electrodes and computing Pearson correlation coefficients between all possible pairs of trials or conditions to form a spatial similarity matrix (L. Wang et al., 2020). Because RSA depends on spatial activity patterns, variations in the number and distribution of electrodes may influence estimated similarity magnitude and, if not carefully interpreted, could affect inferences about representational structure. However, the impact of the number of channel configurations on RSA results has not yet been systematically investigated. This gap directly affects the reliability of conclusions drawn from the growing number of studies especially employing low-density EEG systems.

### 1.1 The aims of the current study

Although recent work has investigated how reducing electrode density affects EEG decoding and has shown that reliable performance can be obtained with substantially fewer channels (Huang et al., 2026). However, these findings come from a narrow set of visually driven paradigms and do not incorporate representational analyses, leaving it unclear whether the effects of channel density generalize across tasks or extend to RSA.

To fill this gap, this study aims to systematically evaluate the impact of different electrode channel configurations (19, 32, 64, and 128 channels) on both MVPA and RSA. We collected two datasets and utilized one publicly available dataset. The first collected dataset used a color harmony assessment with multicolor automotive stimuli; the second employed a visual search task manipulating icon design and activation parameters. The public dataset was a dot-probe task containing both depression patients and healthy controls. All datasets underwent identical preprocessing. The original 128-channel data were then down-sampled to 64, 32, and 19-channel configurations based on the international 10-20 system, generating four data versions for each experiment. For each dataset and channel configuration, we computed time-resolved decoding accuracy and quantified the representational similarity of spatial patterns across experimental conditions. Across multiple datasets and task paradigms, the present study systematically examines how reductions in EEG electrode density affect multivariate analyses at different levels of representation. By comparing decoding and representational similarity based approaches across varying channel configurations, we assess whether the influence of electrode density is consistent across tasks or depends on specific cognitive demands and analysis frameworks.

## 2 Method

We analyzed three EEG datasets to assess how electrode density influences decoding performance and RSA. Two datasets were obtained from our previous experiments, and one publicly available dataset MODMA was included (Cai et al., 2022). The following sections briefly describe the stimuli, tasks, recording procedures, and preprocessing steps for each dataset; full details can be found in the original publications. All studies were approved by the Institutional Review Boards of the corresponding institutions, and all participants provided informed consent.

An overview of the analysis pipeline is illustrated in Fig. 1. Briefly, preprocessed EEG data were first reduced to multiple channel-density configurations via electrode subset selection and segmented into epochs (Fig. 1a). These epoched data were then analyzed in two parallel streams: multivariate decoding to evaluate classification performance (Fig. 1b), and RSA to characterize neural representational structure across channel densities (Fig. 1c).

**Fig. 1.**
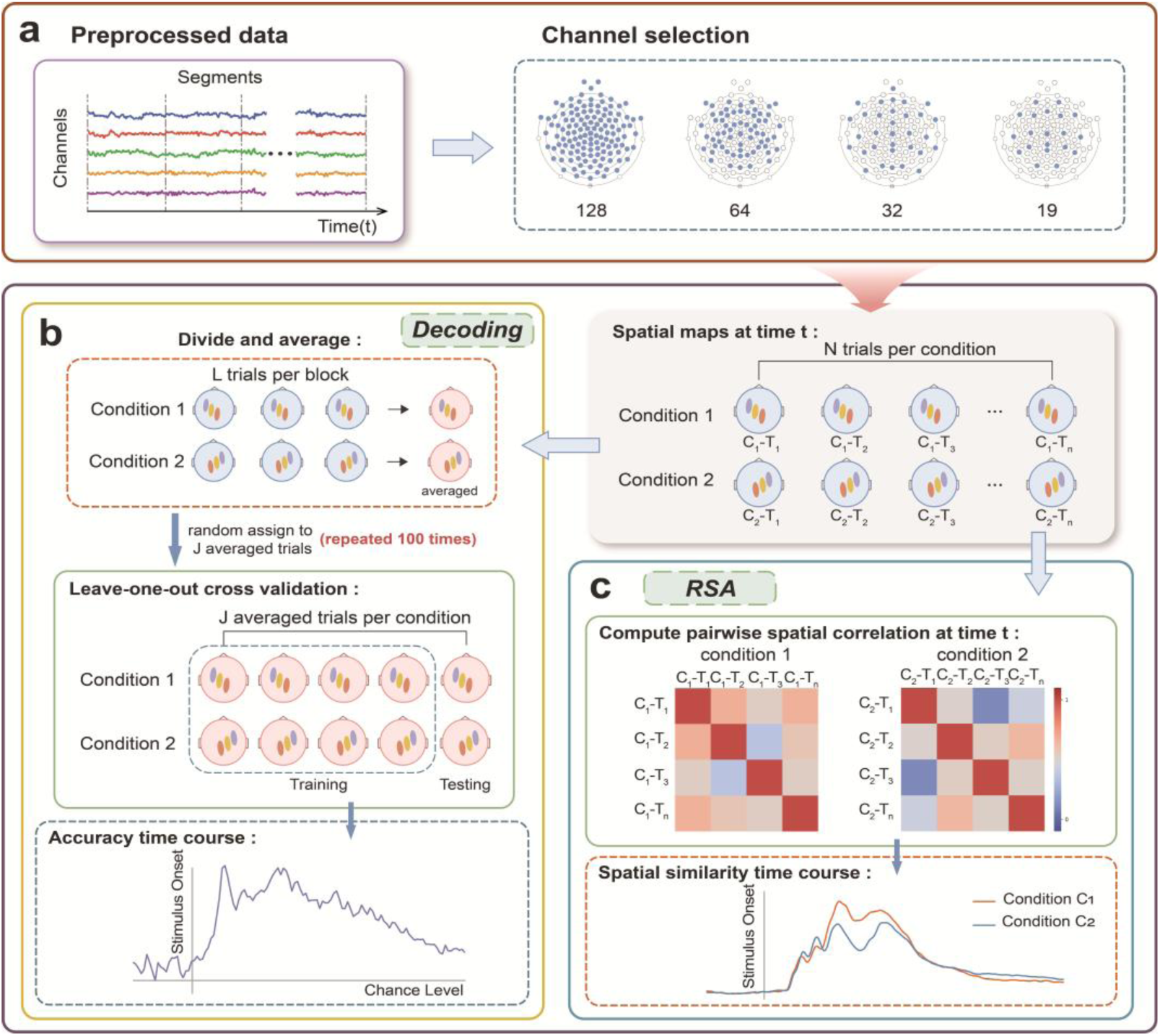
Overview of the analysis pipeline. The workflow consists of three main stages. (a) Preprocessed EEG data were first reduced to four channel-density configurations (128, 64, 32, and 19 channels) via electrode subset selection and then segmented into epochs for subsequent analyses. (b) For each channel configuration, the epoched data were submitted to MVPA to quantify decoding performance. (c) In parallel, the same epoched data were analyzed using RSA to assess how electrode density affects the structure and temporal dynamics of neural representations.

All analyses were conducted in MATLAB 2024a (MathWorks Inc.) using the ERPLAB Toolbox v12.00 (Lopez-Calderon & Luck, 2014). For a straightforward comparison, all datasets underwent identical preprocessing procedures. The original 128-channel EEG data (EGI HydroCel system) were down-sampled to 64-, 32-, and 19-channel configurations by selecting the corresponding electrode subsets defined in the EGI HydroCel montage, which builds on the 10-20 system logic.

### 2.1 Participants

The first experiment included 30 participants (14 males, 16 females; mean age = 29.20 ± 2.38 years; range = 24–34 years), and the second experiment included 30 participants (15 males, 15 females; mean age = 28.87 ± 2.86 years; range = 24–34 years). All participants had normal or corrected-to-normal vision and provided informed consent prior to participation. Both experiments were conducted in accordance with the Declaration of Helsinki and were approved by the Ethics Committee of the University of Jyväskylä (approval ID: l74/13.00.04.00/2024).

The publicly available dataset was obtained from the MODMA database, which includes EEG recordings from 53 participants: 24 outpatients diagnosed with major depressive disorder (MDD; 13 males and 11 females, aged 16–56 years) and 29 healthy controls (20 males and 9 females, aged 18–55 years). After quality control, 29 participants were retained for analysis, comprising 14 patients with MDD (4 males, 10 females; mean age = 27.50 ± 8.47 years) and 15 healthy controls (10 males, 5 females; mean age = 30.07 ± 9.69 years). All participants provided written informed consent prior to participation, and the study protocol was approved by the Ethics Committee of the Second Affiliated Hospital of Lanzhou University, in accordance with the Declaration of Helsinki.

### 2.2 Stimuli and Procedure

#### 2.2.1 Experimental 1: Car Color Harmony EEG Study

This experiment investigated neural responses to color harmony in multicolor automotive designs (Hou et al., 2025). Based on prior design analyses, two-color and three-color vehicle body schemes were created by combining predefined color regions of the car model. Using the Munsell color system and the PCCS-based tone mapping approach, 33 representative colors were selected to form the experimental color palette. From all possible combinations, 100 harmonious and 100 disharmonious color layouts were selected according to the color harmony rule proposed by Moon and Spencer (Moon & Spencer, 1944a, 1944b). The experimental design included two independent factors, namely scheme complexity (two-color versus three-color) and color harmony (harmony versus disharmony), resulting in four experimental conditions: 2C-H, 2C-DH, 3C-H, and 3C-DH. Each condition contained 100 unique stimuli, yielding a total of 400 car images used in the EEG experiment.

The experiment consisted of four condition blocks (2C vs. 3C * H vs. DH) presented in a randomized order. Each block included 100 trials, with image presentation fully randomized. Before the formal experiment, participants completed 20 practice trials (five per condition) to familiarize themselves with the procedure. Each trial began with a 1000 ms fixation cross, followed by a 1000 ms stimulus presentation and a 500 ms blank screen. Participants then rated the valence of each car design on a 7-point Likert scale (1 = very unpleasant, 7 = very pleasant). The scale remained visible until a response was made (Fig. 2a).

**Fig. 2.**
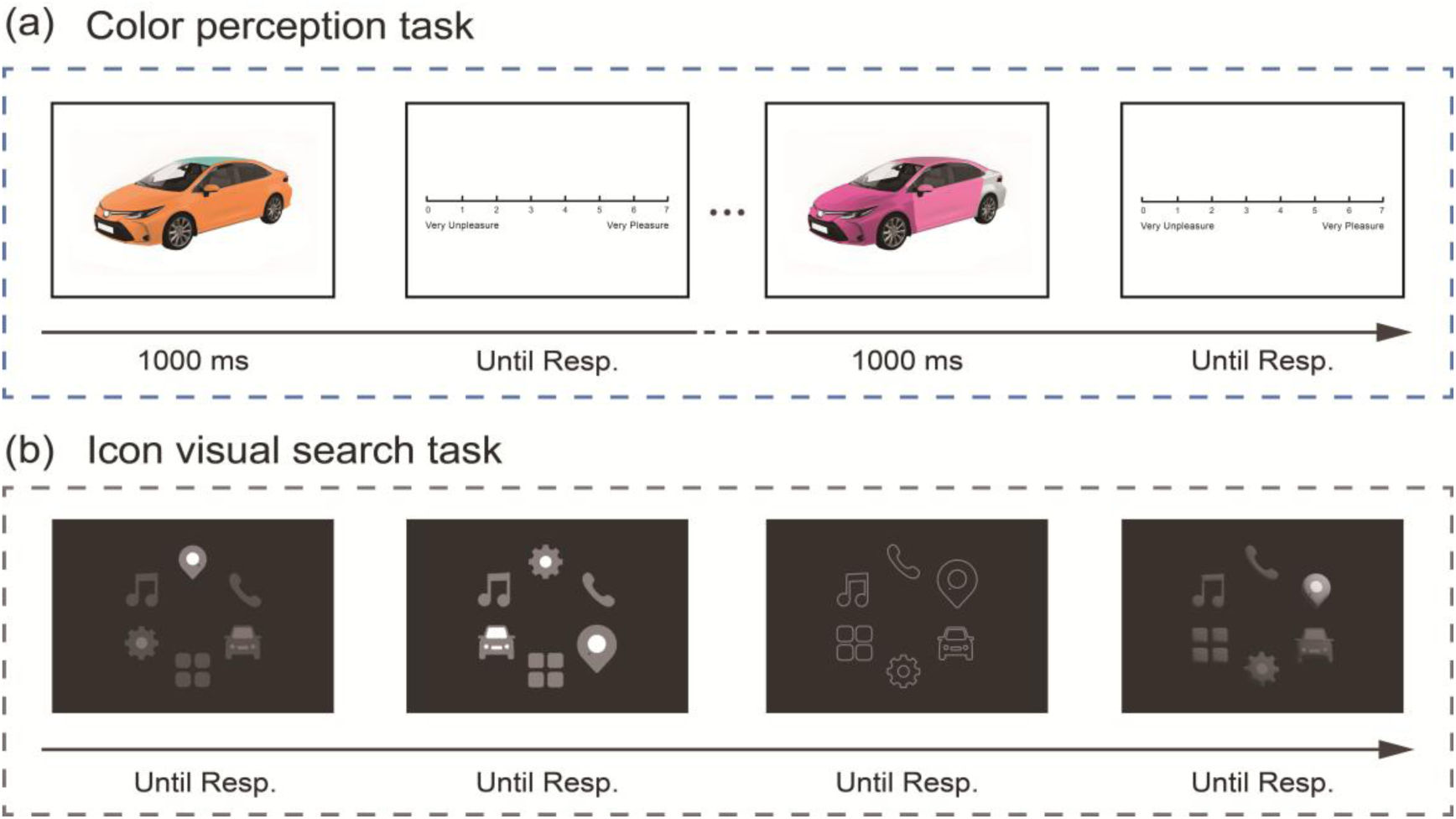
(a) Example stimuli used in the experiment. The figure shows one two-color and one three-color car design. After viewing each image, participants were asked to rate their level of pleasantness on a 7-point Likert scale (1 = very unpleasant, 7 = very pleasant). (b) Example stimuli illustrating the four combinations of icon style and activation mode: flat-highlighted, flat-magnified, linear-magnified, and skeuomorphic-highlighted. Participants were instructed to locate the target icon and judge the position of a small dot embedded within it, pressing “F” if the dot appeared on the left and “J” if it appeared on the right. The stimulus remained on the screen until a response was made.

#### 2.2.2 Experimental 2: Navigation Icon Visual Search Task

This experiment examined the effects of icon style and activation mode on visual recognition using a visual search paradigm (Hou et al., 2026). Based on prior classification, five functional icons were designed in three visual styles (linear, flat, and skeuomorphic) using Adobe Illustrator. Each style was combined with two activation modes (magnified and highlighted), producing six types of stimuli: Lin-Mag (linear magnified), Lin-Hig (linear highlighted), Fla-Mag (flat magnified), Fla-Hig (flat highlighted), Ske-Mag (skeuomorphic magnified), and Ske-Hig (skeuomorphic highlighted).

In each trial, six icons were presented simultaneously on the screen, with the “Navigation” icon serving as the target (see Fig. 2b). The activation mode of the target alternated between activated and non-activated conditions across trials. The remaining five icons served as distractors, sharing the same style and non-activated state. Because correct recognition could not be inferred directly from participants’ visual search behavior, a small dot was embedded within the Navigation icon, positioned either to the left or to the right. Participants were instructed to locate the target icon and indicate the dot’s position by pressing the corresponding key (“F” for left and “J” for right). This design minimized guessing and ensured that participants accurately identified the target icon before responding.

#### 2.2.3 Experimental 3: MODMA Dot-Probe Task

The third dataset was obtained from the publicly available MODMA dataset, which includes EEG recordings from patients with major depressive disorder (MDD) and healthy controls (HC). Further details of the experimental stimuli and procedure are described in the original publication (Cai et al., 2022). The dot-probe paradigm was used to investigate attentional bias toward emotional information. In each trial, participants were asked to maintain fixation on a central cross while viewing pairs of facial images presented side by side on the screen. Each pair consisted of one emotional face and one neutral face, forming three emotional conditions: fear-neutral, sad-neutral, and happy-neutral.

After a short presentation interval, a small white dot appeared randomly on either the left or right side of the fixation point, replacing one of the previously shown faces. Participants were instructed to respond as quickly and accurately as possible to the location of the dot by pressing the corresponding key. This task design allowed assessment of attentional shifts toward emotional stimuli through reaction time and neural responses.

### 2.3 EEG data analysis

#### 2.3.1 Preprocessing

All EEG datasets underwent identical preprocessing procedures. First, a notch filter was applied to remove 50 Hz line noise. The data were then band-pass filtered using a 0.1 Hz high-pass finite impulse response (FIR) filter and a 30 Hz low-pass FIR filter, following EEGLAB’s default zero-phase, Hamming-windowed settings (Luck, 2014; Zhang, Garrett, & Luck, 2024). Subsequently, the data were resampled to 500 Hz and re-referenced to the common average reference, excluding the electrooculogram (EOG) channels (channels 8, 25, 126, and 127) (Jia & Tyler, 2019). Independent component analysis (ICA) was performed using the ICASSO toolbox to identify and remove components associated with ocular and movement artifacts (Zhang, Garrett, Simmons, et al., 2024). Bad channels were interpolated using spherical spline interpolation as implemented in EEGLAB.

Following artifact correction, the EEG data were segmented from 200 ms before to 1000 ms after stimulus onset, except for the car color perception dataset, which was segmented from 200 ms before to 800 ms after stimulus onset. Baseline correction was applied by subtracting the mean voltage of the 200 ms pre-stimulus period from each epoch. Epochs containing extreme amplitudes were identified using ERPLAB’s ‘pop_artmwppth’ moving-window peak-to-peak procedure and subsequently removed. For the car color perception dataset, trials exceeding ±80 µV were discarded, resulting in 91.85% ± 6.01% of trials retained per participant. For the navigation icon judgment dataset, an amplitude threshold of ±100 µV was applied, with 90.30% ± 3.84% of trials retained. For the MODMA dataset, trials exceeding ±100 µV were excluded, resulting in 95.18% ± 5.02% of trials retained per participant.

#### 2.3.2 Multivariate pattern analysis

Time-resolved MVPA was performed separately for each time point for each participant and for each electrode density configuration. All data were resampled to 100 Hz prior to analysis.

Decoding analyses were conducted according to the factorial design of each dataset. For the Color dataset [two within-subject factors design: 2 color harmony (harmony vs. disharmony) × 2 scheme complexity (two-color vs. three-color)], four-way classification was performed across the four experimental conditions (2C-H, 2C-DH, 3C-H, and 3C-DH). In addition, two binary classifications were conducted for each factor: harmony versus disharmony (collapsed across scheme complexity levels) and two-color versus three-color (collapsed across color harmony levels). For the Icon dataset [two within-subject factors design: 3 icon style (linear vs. flat vs. skeuomorphic) × 2 activation mode (magnified vs. highlighted)], six-way classification was performed across all icon conditions. A three-way classification examined differences among icon styles (linear, flat, and skeuomorphic, collapsed across activation modes), and a binary classification assessed activation mode (magnified versus highlighted, collapsed across icon styles). For the MODMA dataset [two within-subject factors design [3 (emotion type: happy vs. sad vs. fear) × 2 (congruency: congruent vs. incongruent)], with two participant groups (healthy controls and MDD),], six-way classification was conducted across all emotional and spatial congruency conditions, along with a three-way classification for emotion category (happy, sad, and fear, collapsed across congruency) and a binary classification for spatial congruency (congruent versus incongruent, collapsed across emotion type).

To overcome overfitting, we used leave-one-out N cross-validation approach as in the previous studies (Zhang, Carrasco, et al., 2024; Zhang, Xin, et al., 2025; Zhang & Luck, 2025). Specifically, the remaining trials were randomly divided into N subsets, and the single-trial EEG signal within each subset was averaged to create ERPset separately. Each subset contained an average of 10-20 trials, as this range has been shown to yield optimal performance for trial averaging (Zhang, Carrasco, et al., 2024). The number of crossfolds was used for each experiment available in Table 1. For each iteration, we used N–1 averaged ERP sets to train the classifier and used the remaining set for testing. This process was repeated N times so that every ERP set was used once for training and testing.

**Table 1.**
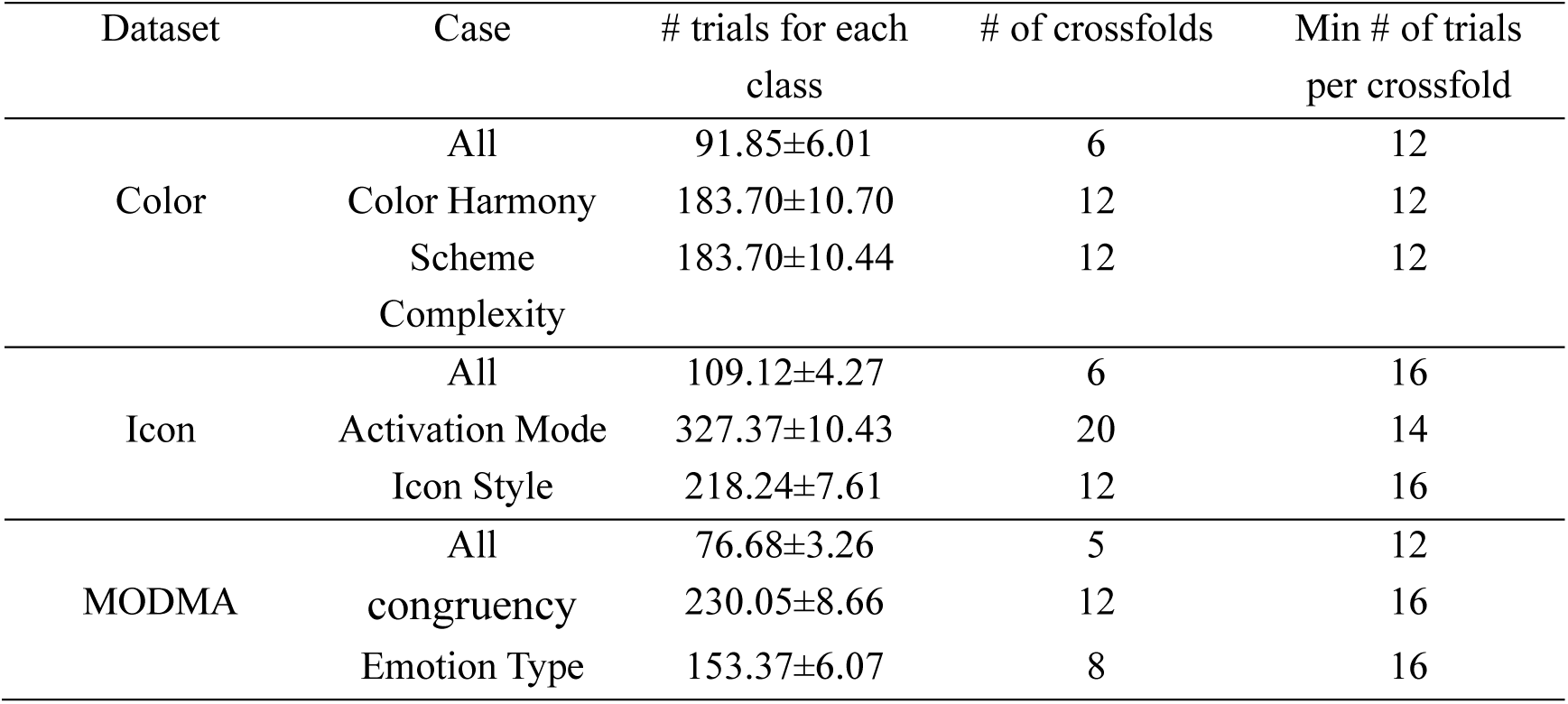
Information about the decoding details for each case and each dataset.

At each time sample, a support vector machine (SVM) classifier was trained and tested. Binary classifications were implemented using the MATLAB function *fitcsvm()*, and multiclass classifications were implemented using *fitcecoc()* with the error-correcting output codes (ECOC) framework (Dietterich & Bakiri, 1994). The MATLAB function *predict()* was used to evaluate classifier performance on one held-out data. To ensure stability of the decoding results, the entire procedure was repeated 100 times. Each participant’s decoding accuracy at each time point was then averaged across the 100 iterations.

#### 2.3.3 Representational similarity analysis

To examine how electrode density influences spatial representational structures, we performed a time-resolved RSA on the three EEG datasets. The same pipeline was conducted to each electrode density configuration.

At each time point, for each participant, a vector representing the spatial pattern of neural activity across all electrodes was extracted. Within each experimental condition, all possible pairs of single-trial vectors were correlated using Pearson correlation coefficient, resulting in a spatial similarity matrix at each time point (Wang et al., 2020). The upper-triangular off-diagonal elements of each matrix were averaged to yield a mean within-condition similarity value. This value indicated the consistency of spatial activation patterns within each condition at one specific time point. This procedure was applied to all datasets and electrode density configurations.

RSA analyses were conducted using the same pipeline as decoding analysis. For the color dataset (2 × 2 design), representational similarity matrices were computed separately for the four experimental conditions (2C-H, 2C-DH, 3C-H, and 3C-DH). To examine the effects of each factor, additional analyses were performed by collapsing across one dimension of the design: data were collapsed across scheme complexity levels to compare harmonious versus disharmonious conditions, and across color harmony levels to compare two-color versus three-color conditions. For the icon dataset (3 × 2 design), representational similarity matrices were computed for all six combinations of icon style (linear, flat, skeuomorphic) and activation mode (magnified, highlighted). To isolate the main effects of each factor, we compared the three icon styles after collapsing across activation modes and contrasted the two activation modes after collapsing across icon styles. Similarly, for the MODMA dataset (3 × 2 design), representational similarity matrices were generated for all six combinations of emotion type (happy, sad, fearful) and position congruency (congruent, incongruent), and follow-up analyses assessed the main effects of emotion (collapsed across congruency) and spatial congruency (collapsed across emotion type).

### 2.4 Statistical analysis

For the decoding results, one-tailed t-tests were first conducted at each time point and for each channel configuration within each dataset to determine whether decoding accuracy significantly exceeded the chance level. P-values were adjusted for multiple comparisons across time points using the false discovery rate (FDR) procedure. We then computed, for each participant and channel configuration, the post-stimulus area under the curve (AUC) of the decoding time course above the chance level to quantify decoding performance. One-way repeated-measures ANOVAs were conducted within each dataset and classification case to test the main effect of channel configuration on decoding performance. Paired sample *t*-tests were then performed to determine whether decoding accuracy under lower electrode densities was significantly reduced compared with that under higher-density configurations. The p values were corrected using FDR.

For the RSA results, the post-stimulus AUC of the RSA time course was computed for each condition and channel configuration. One-way repeated-measures ANOVAs were conducted within each dataset and condition to test the main effect of channel configuration on representational similarity. Paired-sample t-tests were performed for pairwise comparisons across channel configurations, with FDR correction for multiple comparisons.

## 3 Results

### 3.1 Multivariate pattern analysis

To examine the impact of different electrode configurations on decoding performance, we conducted decoding analyses for both all-condition classification and factor-collapsed classifications (i.e., collapsing levels across one factor while decoding the other), separately.

#### 3.1.1 Car color perception dataset

As shown in Fig. 3, decoding accuracies for both the all-condition classification and the binary classification (harmony vs. disharmony and 2-color vs. 3-color) were significantly above chance across nearly the entire post-stimulus time window for all channel configurations. To statistically quantify the channel effects, we computed the post-stimulus AUC of above-chance decoding accuracy at the participant level. A one-way repeated-measures ANOVA revealed no significant main effect of channel configuration for the all-condition classification (F = 1.612, p = 0.190), the color-harmony classification (F = 0.309, p = 0.819), or the scheme-complexity classification (F = 1.052, p = 0.373). We then performed paired sample t-tests across channel configurations, with FDR correction for multiple comparisons.

**Fig. 3.**
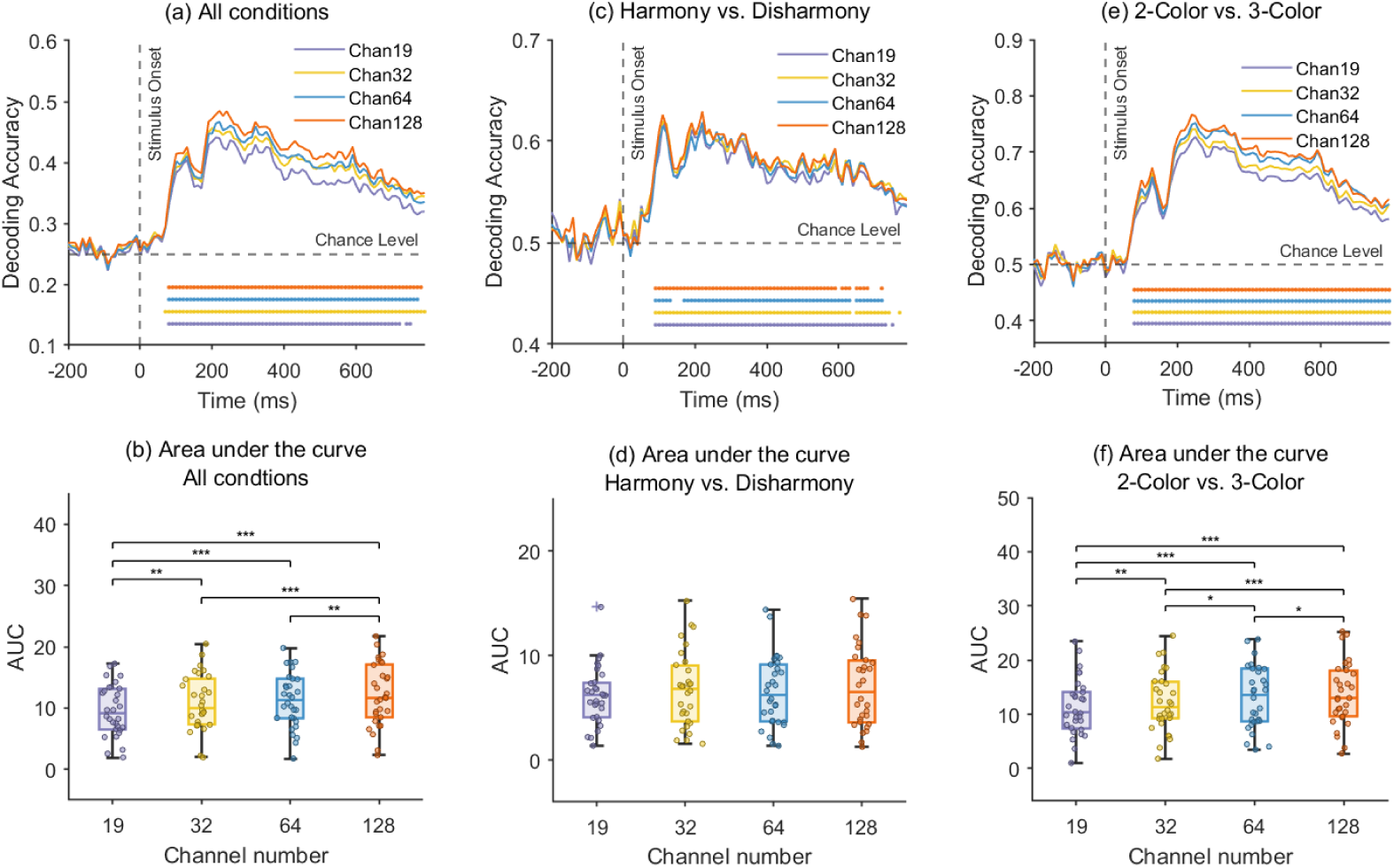
Decoding accuracy results for the Color dataset. The figure summarizes decoding performance across three classification tasks and four EEG channel configurations (16, 32, 64, and 128). (a) Time-resolved decoding accuracies for the all-condition classification, (c) the color-harmony versus color-disharmony classification, and (e) the two-color versus three-color classification, respectively. The horizontal dots mark time points with above-chance decoding accuracy (one-tailed t-tests, FDR corrected). Panels (b), (d), and (f) depict the corresponding post-stimulus AUC of above-chance decoding accuracy for each task. Decoding accuracy was reduced in lower-density configurations in the all-condition and two-color versus three-color classifications, with the 128-channel configuration performing best.

For the all-condition classification (see Fig. 3a and 3b), the post-stimulus AUC of above-chance decoding accuracy for the 128-channel condition was significantly higher than that in the 64-, 32-, and 19-channel configurations (*p* < .001, < .001 and = .001, respectively). The 64- and 32-channel conditions also yielded higher accuracies than the 19-channel condition (*p* = .001 and *p* < .001, respectively). No significant difference was found between the 64- and 32-channel configurations.

For the binary classification of color-harmony (harmony vs. disharmony), no significant differences across channel configurations were observed for the post-stimulus AUC after FDR correction. For the binary classification of scheme complexity (2-color vs. 3-color), decoding accuracies under all channel configurations were significantly above chance throughout the analysis window (see Fig. 3e and 3f). For the post-stimulus AUC of above-chance decoding accuracy, the 128-channel configuration again showed the highest performance, with significant differences relative to the 64-, 32-, and 19-channel configurations (*p* < .001, < .001, and 0.003, respectively). The 64-and 32-channel configurations both outperformed the 19-channel configuration (*p* = 0.008 and <.001, respectively). The 64-channel configuration also outperformed the 32-channel configuration (p = 0.029). For all statistically significant effects observed in Fig. 3, we further quantified the effect sizes by computing the corresponding Cohen’s d values, which are reported in Table 2.

**Table 2.** T statistics and Cohen’s d effect sizes for significant channel-configuration effects in the Color dataset.

| Cohen's $d$ | 32 Vs. 19 | 64 vs. 19 | 128 vs. 19 | 64 vs. 32 | 128 vs. 32 | 128 vs. 64 |
| --- | --- | --- | --- | --- | --- | --- |
| All conditions | 0.294 | 0.372 | 0.550 | 0.071 | 0.266 | 0.202 |
| Scheme complexity | 0.164 | 0.324 | 0.431 | 0.160 | 0.266 | 0.105 |

#### 3.1.2 Navigation icon judgment dataset

Fig. 4 shows the decoding results for the Icon dataset. We conducted three independent analyses: (1) all-condition classification, (2) activation mode classification (magnified vs. highlighted), and (3) icon style classification (linear, flat, and skeuomorphic). Across all three analyses, decoding accuracies for all channel configurations were significantly above the chance level across nearly the entire post-stimulus time window. In general, the average decoding accuracies across channel configurations showed only minor overall differences visually.

**Fig. 4.**
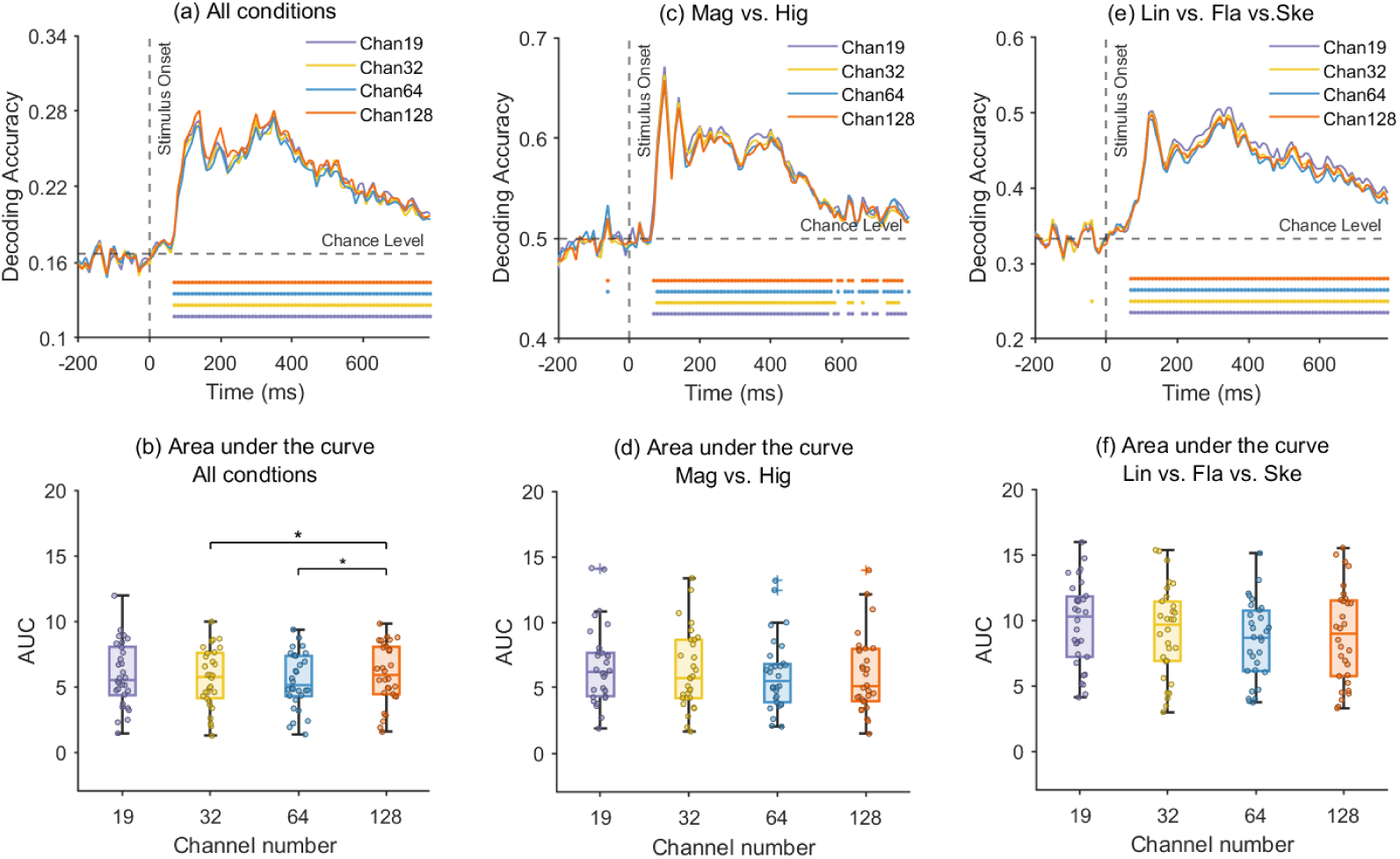
Decoding performance across three classification analyses and four EEG channel configurations (16, 32, 64, and 128). Time-resolved decoding accuracies for (a) the all-condition classification, (c) the activation mode classification (magnified vs. highlighted), and (e) the icon style classification (linear, flat, and skeuomorphic). The horizontal dots denote time points with above-chance decoding accuracy (one-tailed t-tests, FDR corrected). Panels (b), (d), and (f) depict the corresponding post-stimulus AUC of above-chance decoding accuracy for each task. Overall performance was largely stable across different electrode densities, with a slight advantage observed for the 128-channel configuration in the all-condition classification.

To statistically quantify the channel effects, we computed the post-stimulus AUC of above-chance decoding accuracy at the participant level. A one-way repeated-measures ANOVA revealed no significant main effect of channel configuration for the all-condition classification (F = 0.329, p = 0.804), the activation mode classification (F = 0.361, p = 0.781), or the icon style classification (F = 0.765, p = 0.516). We then performed paired sample t-tests (FDR corrected) to examine pairwise contrasts across channel configurations. The results revealed no significant differences among the four channel configurations for the activation mode and icon style classifications. In contrast, for the all-condition classification (see Fig. 4a and 4b), the 128-channel configuration yielded significantly higher mean decoding accuracy than the 64- and 32-channel configurations (p = 0.011, Cohen’s d = 0.225; p = 0.032, Cohen’s d = 0.153, respectively).

#### 3.1.3 MODMA dataset

As shown in Fig. 5, to further assess group-specific decoding performance, classification analyses were conducted separately for the MDD and HC groups. Three types of analyses were performed: (1) all-condition classification, and (2) emotional type classification (happy, sad, and fear). As decoding accuracies for the dot-location classification did not exceed the corresponding chance level under any channel configuration, and no significant differences were observed among the channel conditions, only the results of the all-condition and emotional type classifications are presented.

**Fig. 5.**
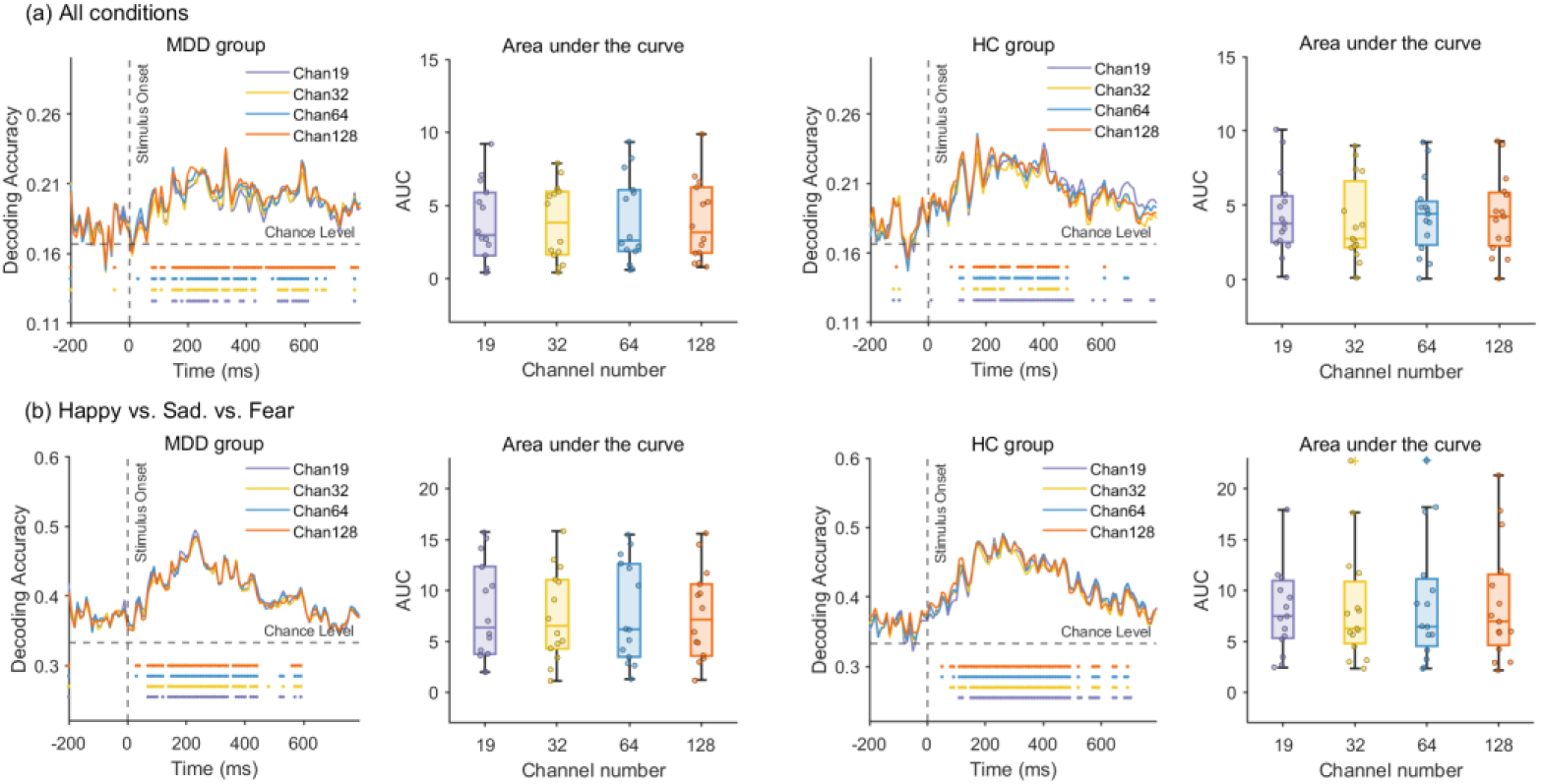
Decoding performance for the MDD Dot-Probe Dataset across four EEG channel configurations (16, 32, 64, and 128) and two groups (MDD and HC). Time-resolved decoding accuracies for (a) the all-condition classification and (b) the emotional-type classification (happy, sad, and fear), and the corresponding post-stimulus AUC of above-chance decoding accuracy for each channel configuration. The dots along the time axis indicate time points where decoding accuracy was significantly above the chance level (one-tailed t-tests, FDR corrected). Both groups showed above-chance decoding across configurations.

For both the all-condition and emotional-type analyses, decoding accuracies across all channel configurations were significantly above chance over nearly the entire post-stimulus interval. To statistically quantify channel effects within each group, we computed the participant-level post-stimulus AUC of above-chance decoding accuracy. A one-way repeated-measures ANOVA revealed no significant main effect of channel configuration for either group in the all-condition classification (HC: F = 0.087, p = 0.967; MDD: F = 0.015, p = 0.998) or in the emotional-type classification (HC: F = 0.029, p = 0.993; MDD: F = 0.024, p = 0.995). We then conducted paired sample t-tests paired (FDR corrected) to examine pairwise contrasts across channel configurations. After correction, none of the pairwise contrasts reached statistical significance in either group.

### 3.2 Representational similarity analysis

To assess whether reducing EEG electrode density alters the spatial representational structure of neural activity, we conducted time-resolved representational similarity analysis (RSA) for all three datasets under four channel configurations (128, 64, 32, and 19 channels). RSA was computed separately for each experimental condition. For statistical comparisons, we quantified representational similarity using the post-stimulus AUC of the RSA time course at the participant level.

#### 3.2.1 Car color perception dataset

Fig. 6 shows the group-averaged temporal dynamics of spatial similarity for the color harmony dataset under different channel configurations. Separate panels depict the four conditions (Harmony, Disharmony, 2-Color, and 3-Color), with each panel comparing RSA time courses obtained from the four electrode densities. Across all four conditions, RSA time courses exhibited broadly similar temporal trends across the four channel configurations throughout the post-stimulus interval.

**Fig. 6.**
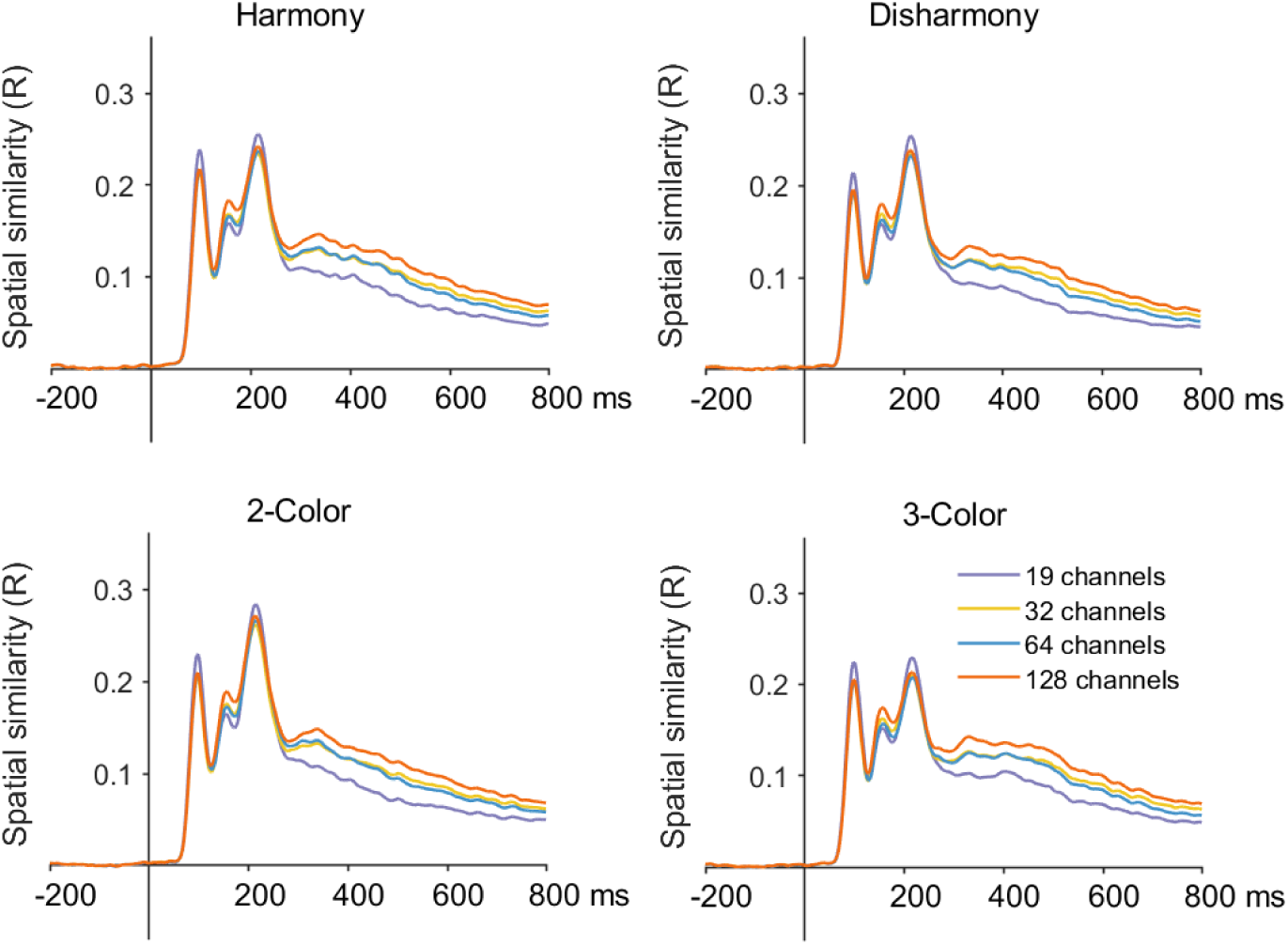
Spatial similarity results for the Color Harmony Dataset across four channel configurations (128, 64, 32, and 19).

To statistically quantify channel effects, we computed the post-stimulus area under the RSA time course (AUC) at the participant level for each condition. A one-way repeated-measures ANOVA revealed no significant main effect of channel configuration for the Harmony condition (F = 1.242, p = 0.298), the Disharmony condition (F = 1.289, p = 0.282), the 2-Color condition (F = 1.267, p = 0.289), or the 3-Color condition (F = 1.156, p = 0.330). We then conducted post-hoc pairwise comparisons using paired t-tests with FDR correction. The corresponding effect sizes (Cohen’s d) for pairwise contrasts are summarized in Table 3. Overall, the temporal evolution of RSA and the relative ordering of conditions were largely preserved across channel configurations.

**Table 3.** Cohen’s *d* effect sizes for spatial similarity contrasts in the Color dataset.

| Cohen's $d$ | 32 Vs. 19 | 64 vs. 19 | 128 vs. 19 | 64 vs. 32 | 128 vs. 32 | 128 vs. 64 |
| --- | --- | --- | --- | --- | --- | --- |
| Harmony | **0.289\*\*** | **0.274\*\*** | **0.512\*\*\*** | -0.034 | **0.201\*\*\*** | **0.246\*\*\*** |
| Disharmony | **0.319\*\*\*** | **0.239\*\*** | **0.513\*\*\*** | -0.090 | **0.176\*\*\*** | **0.275\*\*\*** |
| 2-Color | **0.291\*\*** | **0.287\*\*\*** | **0.513\*\*\*** | -0.023 | **0.202\*\*\*** | **0.236\*\*\*** |
| 3-Color | **0.293\*\*** | **0.236\*\*** | **0.493\*\*\*** | -0.070 | **0.178\*\*\*** | **0.257\*\*\*** |

#### 3.2.2 Navigation icon judgment dataset

Fig. 7 shows the group-averaged temporal dynamics of spatial similarity for the icon dataset across four channel configurations (128, 64, 32, and 19). Separate panels depict RSA time courses for activation mode (Magnified, Highlighted) and icon style (Linear, Flat, Skeuomorphic). Across conditions, RSA time courses exhibited broadly similar temporal trends across the four channel configurations throughout the post-stimulus interval.

**Fig. 7.**
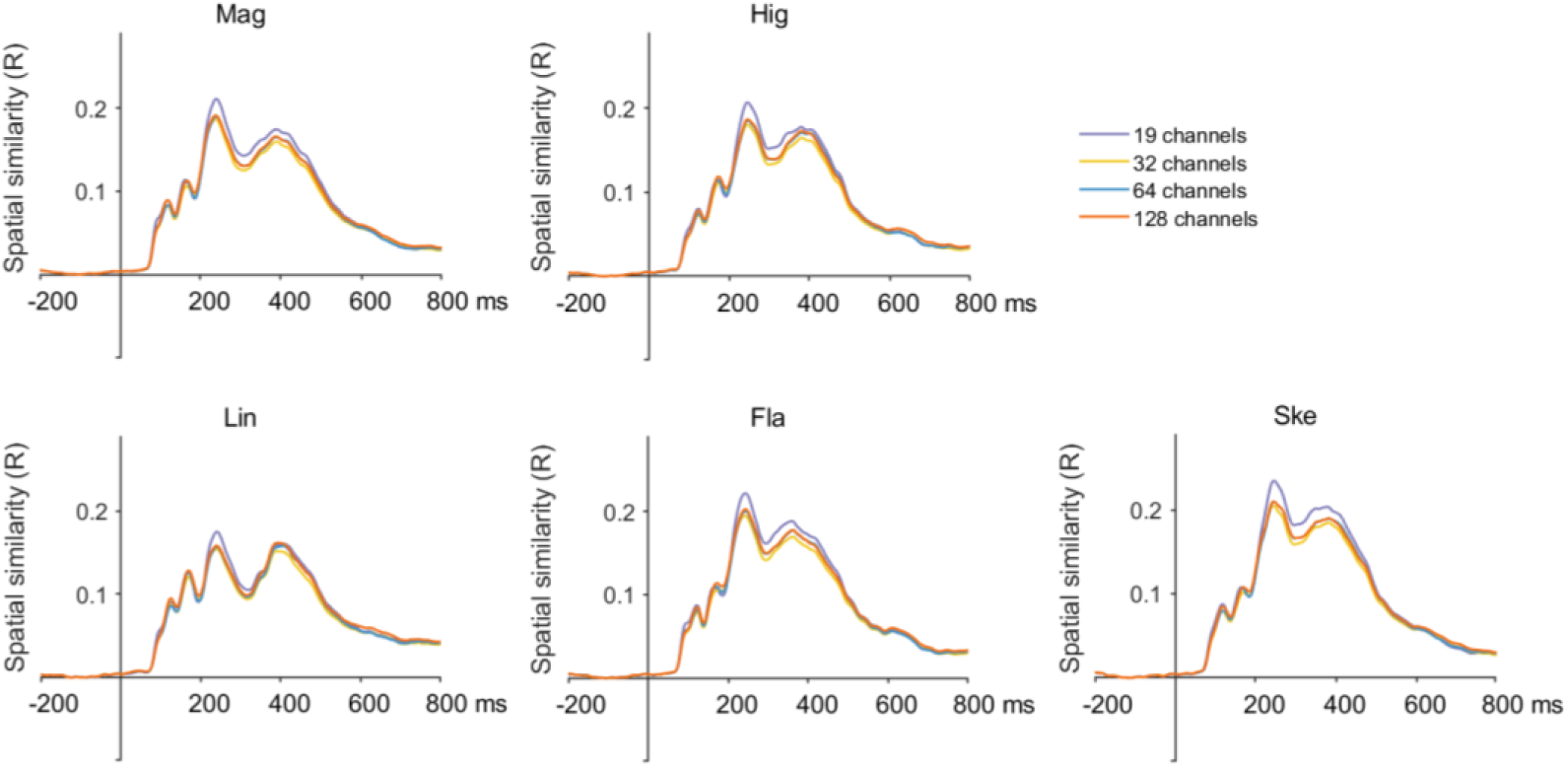
Spatial similarity results for the Icon Design Dataset across four channel configurations (128, 64, 32, and 19).

To statistically quantify channel effects, we computed the post-stimulus area under the RSA time course (AUC) at the participant level for each condition. A one-way repeated-measures ANOVA revealed no significant main effect of channel configuration for the Linear condition (F = 0.285, p = 0.836), the Flat condition (F = 0.281, p = 0.839), the Skeuomorphic condition (F = 0.277, p = 0.842), the Magnified condition (F = 0.288, p = 0.834), and the Highlighted condition (F = 0.303, p = 0.823). We then conducted pairwise comparisons using paired t-tests with FDR correction.

The corresponding effect sizes (Cohen’s d) for pairwise contrasts are summarized in Table 4. Across conditions, the 19-channel configuration generally yielded higher RSA AUC values than the higher-density configurations, whereas differences among the 32-, 64-, and 128-channel configurations were comparatively small.

**Table 4.**
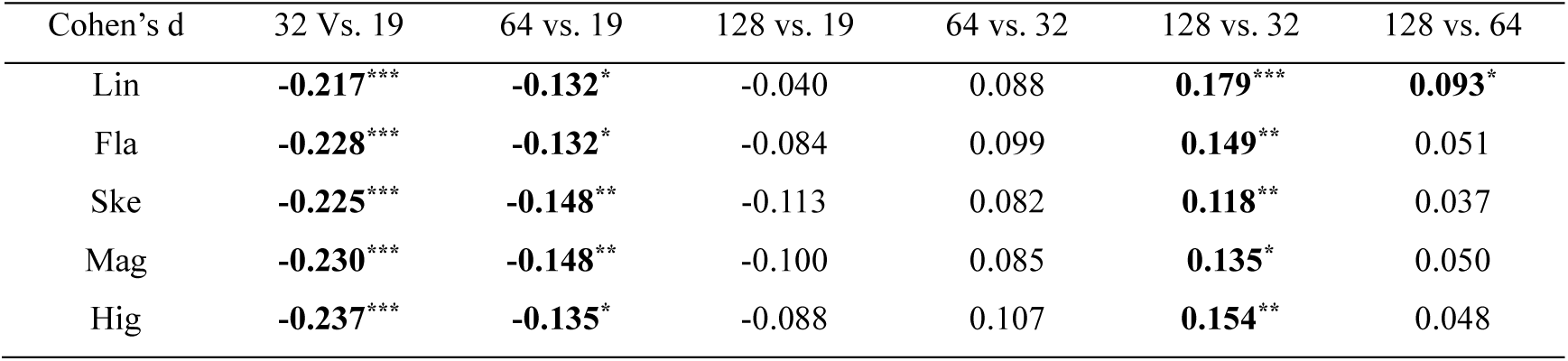
Cohen’s *d* effect sizes for spatial similarity contrasts in the Icon dataset.

| Cohen's $d$ | 32 Vs. 19 | 64 vs. 19 | 128 vs. 19 | 64 vs. 32 | 128 vs. 32 | 128 vs. 64 |
| --- | --- | --- | --- | --- | --- | --- |
| Lin | **-0.217\*\*\*** | **-0.132\*** | -0.040 | 0.088 | **0.179\*\*\*** | **0.093\*** |
| Fla | **-0.228\*\*\*** | **-0.132\*** | -0.084 | 0.099 | **0.149\*\*** | 0.051 |
| Ske | **-0.225\*\*\*** | **-0.148\*\*** | -0.113 | 0.082 | **0.118\*\*** | 0.037 |
| Mag | **-0.230\*\*\*** | **-0.148\*\*** | -0.100 | 0.085 | **0.135\*** | 0.050 |
| Hig | **-0.237\*\*\*** | **-0.135\*** | -0.088 | 0.107 | **0.154\*\*** | 0.048 |

Overall, although absolute similarity values varied with electrode density, most prominently for the lowest-density configuration, the overall temporal profile of RSA was preserved across channel configurations, and the results obtained with the 64- and 128-channel configurations were highly consistent.

#### 3.2.3 MODMA dataset

Fig. 8 shows the group-averaged temporal dynamics of spatial similarity for the MODMA dataset under different channel configurations (128, 64, 32, and 19), plotted separately for the healthy control (HC) and major depressive disorder (MDD) groups and for each emotion condition (happy, sad, and fear). As shown in Fig. 8, RSA time courses obtained from different channel configurations largely overlapped across emotional conditions, with no clear or consistent separation among the four electrode densities.

**Fig. 8.**
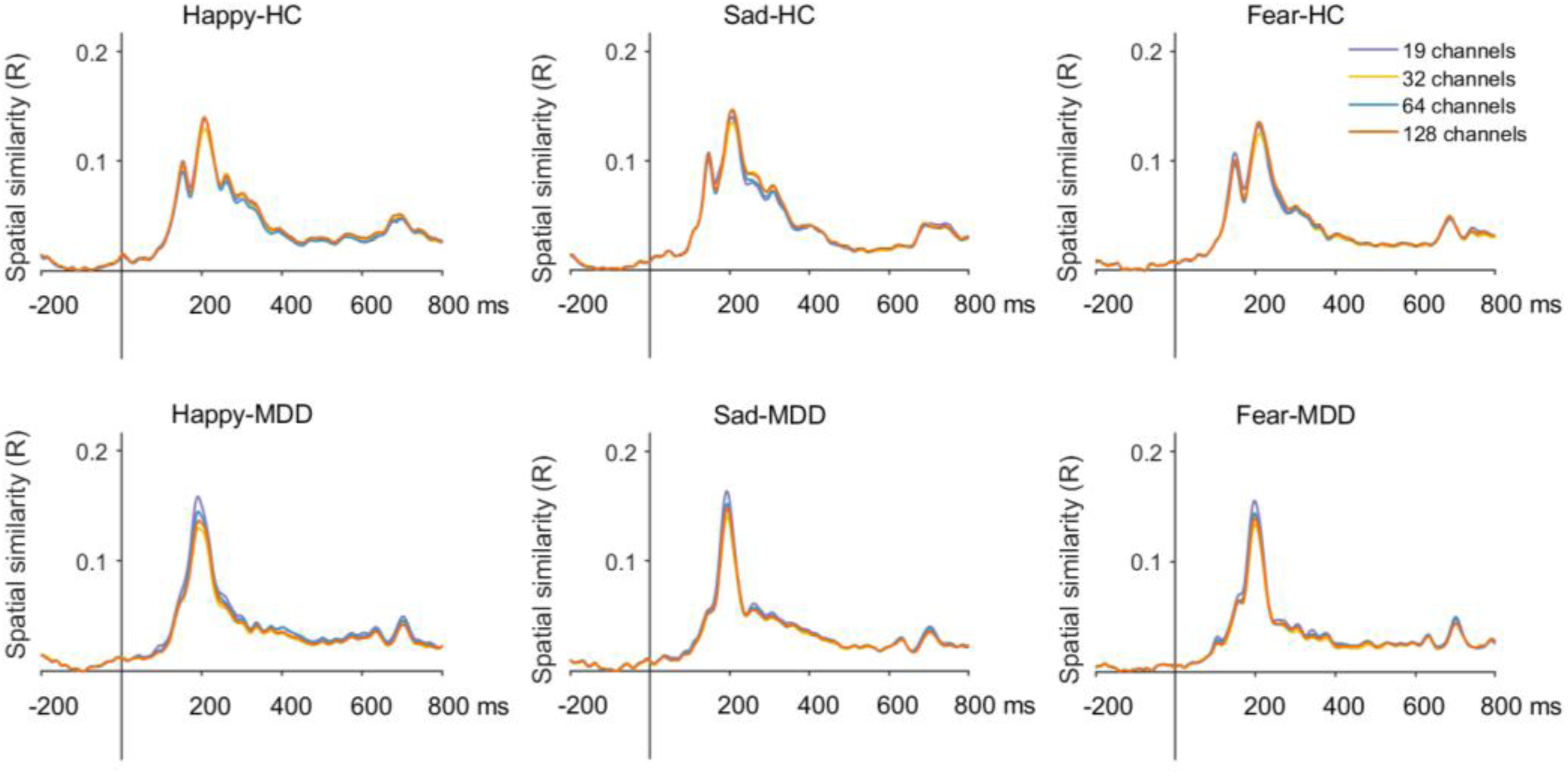
Spatial similarity courses under the happy, sad, and fear face conditions for the HC and MDD groups across four channel configurations (128, 64, 32, and 19).

To statistically quantify channel effects, we computed the post-stimulus area under the RSA time course (AUC) at the participant level for each emotion condition within each group. A one-way repeated-measures ANOVA revealed no significant main effect of channel configuration for the HC group (Happy: F = 0.069, p = 0.977; Sad: F = 0.040, p = 0.989; Fear: F = 0.054, p = 0.983) or for the MDD group (Happy: F = 0.394, p = 0.758; Sad: F = 0.089, p = 0.966; Fear: F = 0.109, p = 0.955). We then conducted pairwise comparisons using paired t-tests with FDR correction, and the corresponding effect sizes (Cohen’s d) are summarized in Table 5. Across all emotions and both groups, RSA time courses were highly similar across the four channel configurations, and no systematic differences were observed.

**Table 5.** Cohen’s *d* effect sizes for spatial similarity contrasts in the MODMA dataset.

| Cohen's $d$ | 32 Vs. 19 | 64 vs. 19 | 128 vs. 19 | 64 vs. 32 | 128 vs. 32 | 128 vs. 64 |
| --- | --- | --- | --- | --- | --- | --- |
| Happy-HC | 0.082 | 0.117 | -0.033 | 0.029 | -0.110 | -0.144 |
| Sad-HC | -0.024 | 0.002 | -0.108 | 0.025 | -0.085 | -0.104 |
| Fear-HC | 0.108 | 0.063 | -0.026 | -0.044 | -0.133 | -0.088 |
| Happy-MDD | **0.347\*** | 0.129 | 0.290 | **-0.229\*** | -0.057 | **0.170\*** |
| Sad-MDD | 0.170 | 0.109 | 0.154 | -0.066 | -0.019 | 0.048 |
| Fear-MDD | 0.206 | 0.061 | 0.122 | -0.138 | -0.079 | 0.059 |

Overall, the MODMA dataset showed minimal sensitivity of spatial representational similarity to electrode density, with largely overlapping RSA time courses across configurations and only isolated, non-systematic differences.

## 4 Discussion

In the current study, we systematically examined how four EEG electrode densities (i.e., 19, 32, 64, and 128 channels) influenced decoding accuracy and spatial representational similarity across three cognitive paradigms. To our knowledge, this is the first study to provide a systematic assessment of how electrode density impacts both decoding sensitivity and representational similarity across distinct paradigms. We found that the effect of electrode density on decoding performance varied across tasks. In tasks where spatial information was more critical (e.g., color complexity classification in the Color dataset), higher electrode densities (64 and 128 channels) significantly improved decoding sensitivity compared to lower densities (19 and 32 channels). However, in other tasks (e.g., icon style classification or emotion decoding in the MODMA dataset), decoding accuracy remained robust even with fewer electrodes, suggesting that some neural representations can be reliably captured with low-density configurations. In contrast, the underlying neural representational structure, as assessed by spatial similarity analysis, was largely consistent across channel densities in terms of relative similarity patterns, indicating that the qualitative organization of neural representations is generally preserved despite variations in electrode count. Together, these results suggest that increasing electrode density selectively enhances decoding sensitivity in a task-dependent manner, while the representational structure remains stable across a wide range of channel configurations. This supports the notion that decoding and representational similarity capture complementary aspects of multivariate EEG signals (Kriegeskorte et al., 2008).

Across the three datasets, decoding performance generally tended to improve with increasing channel count, although the magnitude and statistical significance of these gains were task dependent. Substantial improvements were observed only in the Color Harmony dataset, where decoding accuracy increased noticeably with channel count, likely reflecting finer spatial sampling and more accurate reconstruction of scalp electric fields afforded by higher-density electrode arrays, which facilitate the capture of weaker or more spatially distributed neural activity (Robinson et al., 2017). This pattern suggests that complex perceptual representations such as color combinations rely on spatial information distributed broadly across electrodes, consistent with accounts proposing that visual processing depends on widely distributed cortical networks (Felleman & Van Essen, 1991; Haxby et al., 2001). In contrast, decoding accuracy in the Icon Search and Dot-probe datasets was comparatively stable across channel configurations, indicating that the neural information supporting these tasks is less dependent on dense spatial sampling at the scalp level. For the icon decoding in all conditions, the 128-channel configuration showed small advantages over 64 and 32 channels in the all-condition classification, yet this benefit did not extend to comparisons with 19 channels or to the binary/three-class analyses, suggesting limited incremental gain from additional channels in this task. In the dot-probe dataset, decoding results were broadly comparable across all channel counts, indicating minimal dependence on electrode density under the present task demands. This dissociation highlights that the contribution of electrode density varies with task demands, being more pronounced when the underlying neural representations are spatially distributed and less evident when discriminative information is redundantly expressed or physiologically robust. When decoding performance was quantified by post-stimulus AUC, the one-way repeated-measures ANOVAs did not reveal pronounced main effects of channel configuration across decoding analyses. Nevertheless, pairwise contrasts revealed statistically reliable density-related advantages in the Color dataset, particularly for the all-condition and scheme-complexity classifications, whereas channel-configuration effects were limited in the Icon and MODMA datasets.

Despite task-related differences, decoding performance remained significantly above chance even under the 32- and 19-channel configurations in all three datasets. Notably, the magnitude of channel-related effects was highly task-dependent: whereas the Color dataset showed clear benefits of higher-density configurations, effect sizes in the Icon and MODMA datasets were small or negligible, indicating limited gains from additional electrodes when decoding performance was already robust across channel counts. These findings suggest that, at least for the cognitive processes examined here, discriminative neural information exhibits a degree of spatial redundancy, such that reliable classification can be achieved even with sparse spatial sampling. Although high-density EEG improves spatial resolution and can enhance decoding sensitivity, our results show that low-density configurations can still support reliable decoding (Huang et al., 2026; Xi et al., 2025), particularly when the underlying neural signatures are strongly expressed or redundantly distributed. This pattern aligns with prior work showing that increasing electrode count improves classification performance within low-to-moderate density ranges (Robinson et al., 2017), but extends this literature by demonstrating that such improvements are not universal and may plateau or diminish depending on task demands and representational properties.

Whereas decoding performance showed clear channel-dependent effects that varied across tasks, the RSA analyses exhibited a different pattern, with the relative representational structure remaining largely stable across electrode configurations. RSA captures the relative similarity structure between conditions and therefore exhibits greater robustness under moderate reductions in spatial sampling (Kriegeskorte et al., 2008). It should be noted that the higher absolute RSA values observed under the 19-channel configuration do not necessarily indicate superior neural representational quality. Instead, this pattern may be related to the sensitivity of correlation-based similarity measures to the dimensionality of spatial activation patterns. RSA relies on correlations between spatial vectors across electrodes, and when fewer channels are included in the analysis, the dimensionality of these vectors is reduced. This reduction may lead to a compression of variance across conditions, thereby increasing the likelihood of obtaining higher correlation coefficients. Consequently, elevated RSA values observed under low-density configurations are more likely to reflect a bias in correlation estimates introduced by channel reduction, rather than necessarily a genuine enhancement of neural representations. Despite differences in absolute similarity magnitude across channel configurations, the direction of condition differences and the corresponding effect sizes remained consistent. These findings suggest that task-relevant representational structure is robust and reproducible across electrode densities, even when absolute RSA values vary. Importantly, the one-way repeated-measures ANOVAs based on AUC measures revealed no significant main effects of channel configuration for RSA across datasets and experimental conditions, further supporting the robustness of representational similarity structure to variations in electrode density.

Once the critical sources of information are effectively captured, further increases in electrode number may primarily introduce spatial redundancy, rather than fundamentally altering the representational geometry (Grootswagers et al., 2017). Consistent with this view, our results showed that although reducing electrode density modestly weakened the fine-grained stability of RSA patterns in tasks with higher perceptual complexity, the overall relative structure of representational similarity was largely preserved across channel configurations. This interpretation is strengthened by effect size analyses, which revealed that RSA effect magnitudes were highly consistent across electrode densities: whether representational differences were present or absent, the direction and size of effects showed minimal change with channel reduction, highlighting the robustness of RSA to variations in spatial sampling. Nevertheless, in conditions requiring greater perceptual integration (e.g., color-harmony decoding), subtle representational distinctions became less pronounced at lower densities, indicating that spatial sampling benefits are task-dependent rather than universal. Taken together, these findings suggest that low-density EEG is generally sufficient to characterize representational structure at the level of condition similarity, while high-density systems retain clear advantages when analyses demand finer spatial precision or when representational differences are subtle and spatially distributed.

Several limitations should be considered. First, our conclusions regarding the differential effects of channel count are based on comparisons among four discrete electrode configurations. Although these configurations span a representative range, examining intermediate densities or identifying optimal electrode subsets could yield more refined insights into the saturation point of decoding performance and the minimum number of channels required for stable RSA. Second, the generalizability of the present findings remains to be established in broader populations and recording contexts, such as developmental or aging cohorts and alternative EEG acquisition systems (e.g., dry electrodes) (Kam et al., 2019). Although the three datasets analyzed here are diverse, they primarily involve visual processing, and future work should extend these investigations to other cognitive functions. Third, the current conclusion from linear SVM-based decoding analysis may not generalize to the other advanced algorithms, convolutional neural networks, recurrent neural networks.

## 5 Conclusion

This study provides a systematic cross-paradigm evaluation of how EEG electrode density shapes multivariate inferences from scalp recordings. By integrating MVPA and RSA within a common preprocessing and analysis framework, we establish a clear distinction between sensitivity and structure: reducing channel count primarily affects the sensitivity of multivariate readouts, whereas the qualitative organization of representational relationships remains largely preserved across a wide range of montages. Importantly, this pattern holds across cognitive domains, indicating that electrode density is a critical design parameter for maximizing effect size and reliability, but not necessarily a prerequisite for obtaining meaningful decoding and representational results.

Overall, the present work offers empirical guidance for selecting EEG channel counts under different spatial and practical constraints. It supports the feasibility of conducting robust multivariate analyses with low-density EEG when high-density systems are impractical, while highlighting the added value of dense arrays when fine-grained representational stability or maximal decoding sensitivity is required. Future work should further establish generalizability to broader populations and recording contexts and examine whether montage- or subject-specific electrode selection can further improve efficiency under low-density configurations.

